# Benchmarking long-context genome language models on biosynthetic gene clusters

**DOI:** 10.64898/2026.05.12.724296

**Authors:** Keisuke Hirota, Koichi Higashi, Ken Kurokawa, Takuji Yamada

## Abstract

Recent advances in language models for natural language processing have spread to the field of genomics, driving the development of genome language models (gLMs) to decipher genomic information. Cutting-edge long-context gLMs are promising approaches for understanding and designing biological complexity, but their evaluation remains underdeveloped. In this study, we introduce BGCs-Bench, a unified benchmark focused on biosynthetic gene clusters for assessing long-range genomic modeling on three downstream tasks: biosynthetic class prediction, taxonomic classification and coding sequence annotation. Using BGCs-Bench, we perform systematic and layer-wise evaluations of the embedding representations of long-context gLMs, demonstrating that layer selection is crucial for downstream task performance. In addition to the evaluation results, the logit lens analysis of autoregressive gLMs suggests that StripedHyena-based models consist of earlier layers to encode biologically meaningful information from input DNA sequences and deeper layers to optimize embeddings for sequence generation. These findings provide insights for more effective development and application of long-context gLMs.

## Introduction

DNA, the language of life, encodes the genetic blueprint and understanding its grammar, syntax and semantics has remained a long-standing challenge in biology. On the basis of the analogy between DNA and language, natural language processing (NLP) techniques have been applied to genomics [1,2]. Deep learning approaches have achieved remarkable success in various genomic tasks such as DNA promoter identification [3,4] and gene annotation [5–8]. In particular, the Transformer architectures [9] and improvements in computer hardware have allowed the paradigm of large language models (LLMs) based on a large-scale pretraining strategy, driving the development of genome language models (gLMs) for decoding genomic information. Transformer-based gLMs including DNABERT [10], DNABERT-2 [11] and Nucleotide Transformer [12] have shown performance comparable to that of task-specific models in diverse downstream tasks; however, these gLMs have limitations in term of context-window size because of the quadratic computational cost of the self-attention mechanism. Previous studies have approached this bottleneck by applying various tokenization strategies [11,13] and alternative architectures [14–16]. HyenaDNA [14] was the first gLM to achieve a 1 Mbp-context with single-nucleotide resolution via the Hyena operator with subquadratic computational cost. Subsequently, Evo [17] and Evo 2 [18] were developed as 131 kbp- and 1 Mbp-context foundational gLMs, respectively, by leveraging StripedHyena, the hybrid architecture combining Hyena and attention layers.

Long-context gLMs substantially broaden the scope of genomic modeling from relatively localized units such as single genes or compact modules such as toxin-antitoxin systems to higher-order genomic organization spanning functional modules, including biosynthetic gene clusters (BGCs). BGCs are representative examples of such long-range functional modules and comprise co-localized genes that collectively encode the biosynthesis of specialized metabolites. Despite the rapid emergence of long-context gLMs, their evaluation remains underdeveloped. Several benchmark efforts have begun to address this gap. BEND [19] introduced a collection of biologically meaningful tasks for gLMs, including long-range downstream tasks such as enhancer annotation. Recently, the genomics long-range benchmark (LRB) [20] and DNALONGBENCH [21] explicitly focused on tasks requiring long-range dependencies, thereby extending evaluations spanning 100 kbp to 1 Mbp scales. These benchmarks highlight that some gLMs capture only limited information about long-range features, but the latest StripedHyena-based gLMs were not included in these studies and therefore require further assessment.

In addition, existing benchmarks have focused primarily on the comparative performance of models and how latent representations of gLMs can be most effectively extracted and utilized in practice remains unclear. This issue is particularly critical for embedding-based downstream analyses because previous studies on language and biological sequence modeling have shown that latent representations are not functionally homogeneous across layers. Regarding NLP, an analysis of BERT demonstrated that different layers encode different aspects of syntactic structure [22]. A similar pattern has been observed in protein language models: the intermediate layers are informative for family- and function-related signals, whereas the final layer is tied to structural signals in ESM-2 [23]. Evo 2 also appears to exhibit layer-wise characteristics; it has been reported that some intermediate layers are suitable for sparse autoencoder-based interpretation and encode a phylogenetic manifold [18]. These observations underscore the need for systematic layer-wise evaluation of long-context gLMs.

Another limitation is that existing benchmarks often rely on different DNA sequence datasets for different tasks [11,12,19–21,24]. While this practice is reasonable for task-specific model development, it complicates cross-task interpretation: observed performance differences may reflect not only differences in model capability but also differences in the underlying datasets. Consequently, disentangling the effects of task type, dataset composition and embedding strategy remains difficult when comparing gLMs.

Taken together, these benchmark limitations in terms of targets, evaluation strategies and datasets indicate that the critical assessment of long-context gLMs remains fragmented. Existing benchmarks have established the importance of long-range genomic modeling, but they provide only a partial view of model behavior. Key factors directly affecting the practical use of gLMs, including layer selection, pooling strategies and the interaction between representation quality and downstream task type have not yet been systematically examined. Therefore, a unified benchmark that explicitly addresses these issues is needed to achieve a more comprehensive understanding of foundational gLMs.

Here, we present BGCs-Bench, a unified benchmark that defines multiple biologically relevant tasks on the same DNA sequence dataset for long-range genomic modeling with a focus on BGCs as the long-range modules encoding multifaceted biological information (Figure 1). Using BGCs-Bench, we perform a consistent and multidimensional evaluation of existing foundational gLMs with a focus on how representation quality varies across model depth, pooling methods and downstream tasks. Our study provides both a novel benchmark resource for long-range DNA prediction tasks and practical insight into the analysis and use of representations learned by long-context gLMs.

**Figure 1.**
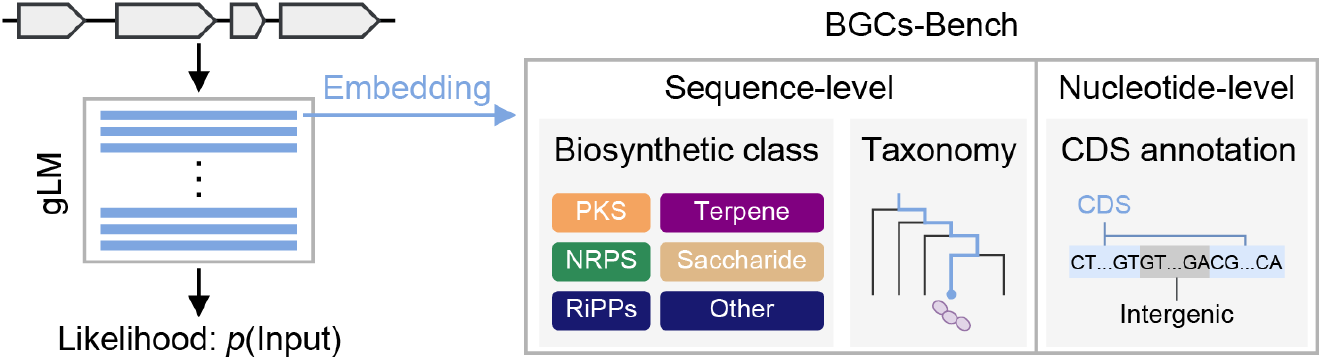
Overview of BGC-Bench. BGCs-Bench contains three downstream tasks to assess the embedding quality of long-context gLMs: biosynthetic class prediction, taxonomic classification and CDS annotation.

## Materials and methods

### BGCs-Bench construction

We downloaded DNA sequences and annotation data of BGCs from MiBiG version 4.0 [25]. To ensure data quality, BGC entries with “retired” or “pending” statuses were excluded, resulting in 2,437 “active” entries that are currently maintained. Given the context length of existing gLMs and computational constraints, we further filtered the dataset by sequence length using thresholds of 131,072 bp and 65,536 bp to enable fair evaluation across models with different practical input length limits caused by the maximum context window size and required computational cost. This procedure yielded two datasets comprising 2,386 and 2,044 BGCs, hereafter referred to as the 131k dataset and the 66k dataset, respectively. To evaluate sequence-level embeddings, each BGC was assigned functional and taxonomic labels on the basis of the MiBiG annotations. The functional labels were defined as biosynthetic classes, a multilabel classification that included six categories: NRPS, PKS, RiPPs, terpene, saccharide and other. Taxonomic labels were assigned at the family level using TaxonKit (v0.20.0) [26]. Both datasets were split into five folds for cross-validation, ensuring a balanced distribution of functional and taxonomic labels across the folds.

In addition, we constructed a subset, hereafter referred to as the annotation dataset, to evaluate nucleotide-level embeddings. For both prokaryotic and eukaryotic BGCs, we sampled 10 sequences from each biosynthetic class that had at least 10 available sequences, excluding “other”, resulting in a total of 80 BGCs. Every nucleotide was labeled as coding sequence (CDS) or intergenic region according to its MiBiG annotation. The total size of the annotation dataset was 2,098,355 bp. The annotation dataset was also divided into five folds with a balance between both biosynthetic class distribution and the proportion of prokaryotic versus eukaryotic sequences across folds.

### Retrieval of gLM embeddings

To evaluate representative autoregressive long-context gLMs, we selected HyenaDNA [14], Evo [17] and Evo 2 (7B and 40B) [18] (Table 1). For each model, layer-wise embeddings of BGC sequences were extracted from all blocks using either the official codebase or PyTorch hook functions. As sequence-level representations, we examined mean-pooling embeddings, which were obtained by averaging nucleotide-level embeddings over the full sequence, and last-token embeddings, which were taken from the final token position. We additionally evaluated the embeddings of reverse-complement sequences to assess the effect of strand orientation. To compare the quality of sequence-level embeddings, we further included NTv3 [27], a U-Net-based long-context gLM, for which only the mean-pooling strategy was considered (Table 1). Inference for Evo 2 was performed on eight NVIDIA H200 GPUs, whereas all the other models were run on a single NVIDIA H200 GPU.

**Table 1.** gLMs evaluated in this study.

| Model | Architecture | Parameter | Context size |
| --- | --- | --- | --- |
| HyenaDNA | Hyena | 54.6 M | 1 Mbp |
| Evo | StripedHyena | 7 B | 131 kbp |
| Evo 2 | StripedHyena2 | 7 B or 40 B | 1 Mbp |
| NTv3 | U-Net | 650 M | 1 Mbp |

### Linear probing

We assessed the quality of the gLM embeddings using a logistic regression classifier (scikit-learn v1.7.0 [28]) as a linear probe for each downstream task: biosynthetic class prediction, taxonomic classification and CDS annotation. To evaluate sequence-level embedding evaluation, we also implemented baselines based on k-mer frequency features from 3-mers to 9-mers. In both the multilabel and multiclass classification settings, classifiers were trained in a one-vs-rest fashion for each class. The hyperparameters of the logistic regression classifiers were set to default values, except that the maximum number of iterations was set to 1,500 for the CDS annotation task and to 1,000 for the other two tasks to ensure convergence, and the weighted loss function based on the class weights was adopted to address the class imbalance. Performance was evaluated primarily using Macro F1 for BGC type and taxonomic classification and Matthew’s correlation coefficient (MCC) for CDS annotation.

In the biosynthetic class prediction task, inclusion of the “other” category was treated as optional, and analyses were conducted both with and without this category. In the taxonomic classification task, family-level prediction was performed on the top 10 families by BGC count, and analyses were likewise conducted both including and excluding the remaining families, including *Unclassified*.

### Logit lens

To further characterize the latent representations of autoregressive gLMs, we employed the logit lens [29]. The logit lens is a training-free interpretability technique for language models, which projects intermediate hidden states directly into the model’s output vocabulary space via the unembedding layer. This method thereby converts embeddings into logits, allowing tracking of how next-token predictions emerge and are progressively refined across layers.

We applied the logit lens to all layers of the autoregressive gLMs: HyenaDNA, Evo, and Evo 2 (7B and 40B). For each layer, the latent representation was passed through the corresponding unembedding layer to obtain logits, which were then evaluated by negative log-likelihood (NLL) using the 66k dataset. This analysis enabled us to assess how predictive information emerges across layers and to characterize the contributions of different depths, especially for large-scale gLMs.

## Results

### Details of BGCs-Bench

BGCs-Bench comprises three datasets designed to evaluate long-context gLMs from complementary perspectives: the 131k dataset, the 66k dataset and the annotation dataset. In the datasets for assessing sequence-level embeddings, the biosynthetic type was dominated by NRPS and PKS, each accounting for approximately 30% of the BGCs (Supplementary Figure S1 and Supplementary Figure S2). With respect to taxonomy information, Streptomycetaceae was by far the most abundant family, representing approximately 30% of each dataset, followed by Aspergillaceae, which accounted for approximately 10% (Supplementary Figure S1 and Supplementary Figure S2). The annotation dataset for assessing nucleotide-level embeddings consisted of 50 prokaryotic and 30 eukaryotic BGCs, corresponding to approximately 1.3 million and 0.8 million nucleotides, respectively (Supplementary Figure S3). Across these 80 BGCs, CDSs outnumbered intergenic regions by 4-fold on average.

### Evaluation of downstream tasks via linear probing

We first evaluated the quality of the sequence-level embeddings via linear probing for two downstream tasks: biosynthetic class prediction and taxonomic classification. The results on the 66k dataset for functional classification revealed that gLMs captured functional signals from both DNA sequences and their reverse complements (Figure 2A and Supplementary Figure S4). Furthermore, mean-pooling embeddings consistently outperformed last-token embeddings for autoregressive models. Compared with the baseline classifiers based on k-mer frequency, HyenaDNA underperformed across all layers, whereas NTv3 consistently outperformed the baseline throughout the depth (Figure 2B). In contrast, the performance of StripedHyena-based gLMs depended strongly on layer selection. This effect was particularly remarkable for Evo 2 (40B), in which some layers surpassed that of other gLMs, while others performed worse. Furthermore, StripedHyena-based gLMs showed a consistent pattern in which the predictive performance decreased after a certain layer (Figure 2C). A marked decrease in performance after specific layers was likewise observed regardless of whether the 66k or 131k dataset was used and whether the “Other” class was included in the classification (Supplementary Figure S5).

**Figure 2.**
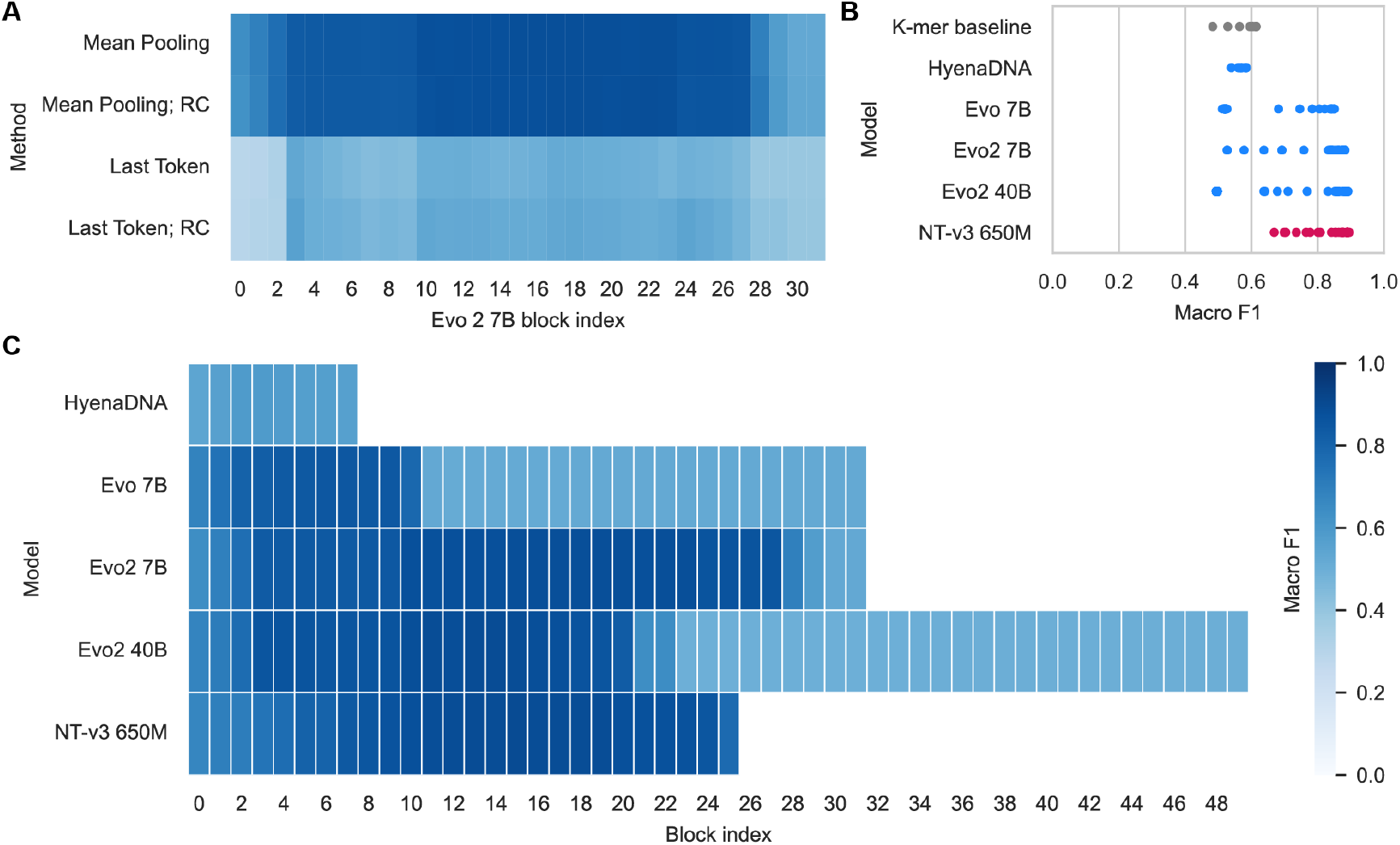
Results of the biosynthetic class prediction. These results show the predictive performance of mean-pooling embedding-based logistic regression classifiers on the 66k dataset excluding the “Other” class. (A) Performance of Evo 2 (7B)-based classifiers. (B) Comparison of k-mer (k=3~9) baselines and blocks of gLMs. (C) Overview of the predictive performance per layer in the gLMs.

We next assessed the taxonomic classification using the same procedure as that used for biosynthetic class prediction. The results on the 66k dataset similarly revealed that gLMs captured taxonomic signals from both DNA sequences and their reverse complements and that mean-pooling embeddings yielded better performance than last-token embeddings did in autoregressive models (Figure 3A and Supplementary Figure S6). All gLMs except HyenaDNA outperformed the k-mer baselines when an appropriate layer was chosen (Figure 3B). With respect to functional classification, StripedHyena-based gLMs consistently exhibited a decrease in predictive performance after a particular layer (Figure 3C). This notable deterioration pattern of gLM embeddings was robust across both the 66k and 131k datasets and regardless of whether the “Other” families were included in the classification (Supplementary Figure S7).

**Figure 3.**
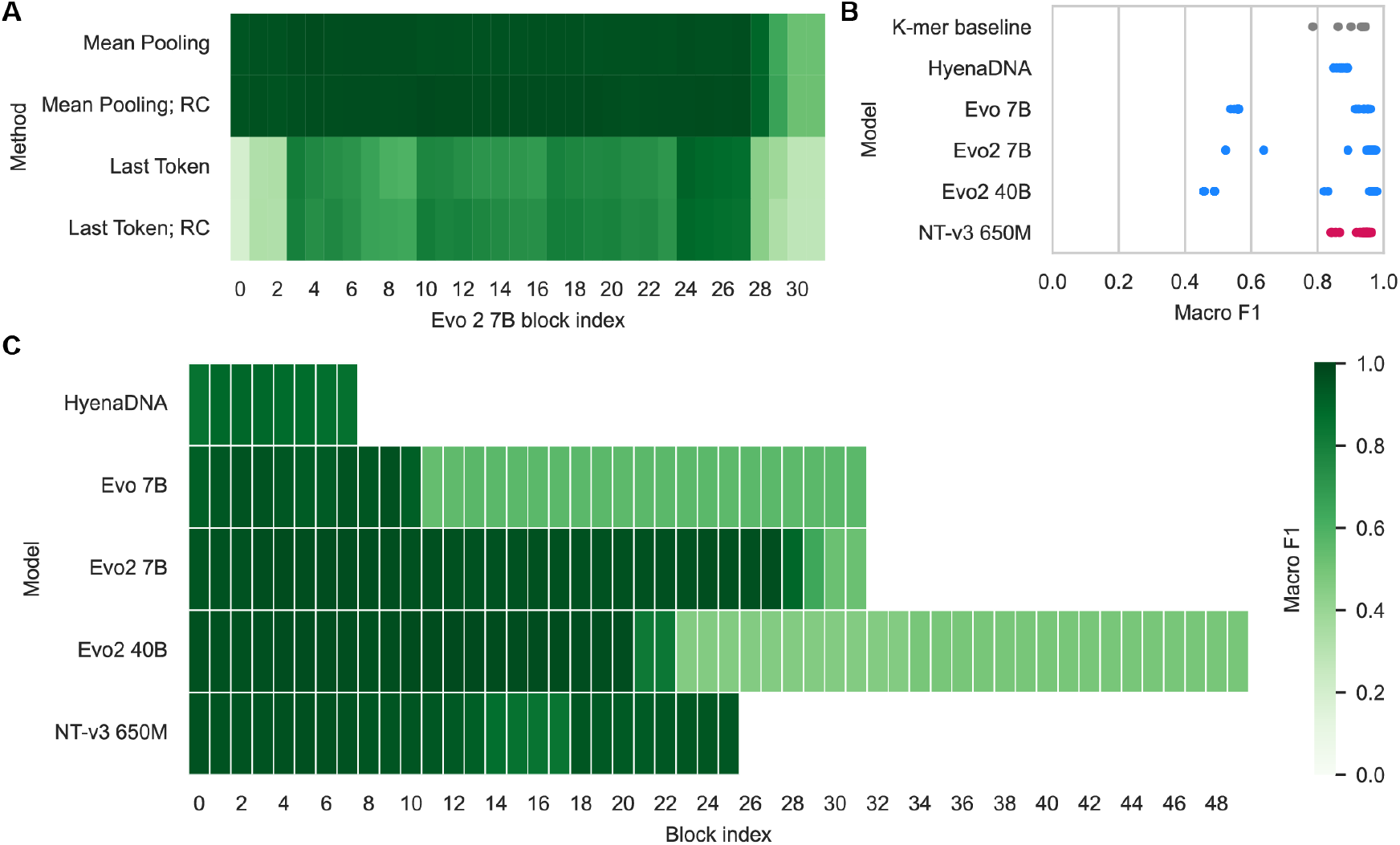
Results of the taxonomic classification. These results show the predictive performance of mean-pooling embedding-based logistic regression classifiers on the 66k dataset excluding “Other” families. (A) Performance of Evo 2 (7B)-based classifiers. (B) Comparison of k-mer (k=3~9) baselines and blocks of gLMs. (C) Overview of the predictive performance per layer in the gLMs.

Finally, we examined the quality of the nucleotide-level embeddings via linear probing on the CDS annotation task. Similar to the results for sequence-level embeddings, StripedHyena-based gLMs achieved strong performance at specific layers but consistently exhibited a decrease in performance beyond a certain depth (Figure 4). For the Evo and Evo 2 models, the depth at which the collapse emerged was consistent across downstream tasks.

**Figure 4.**
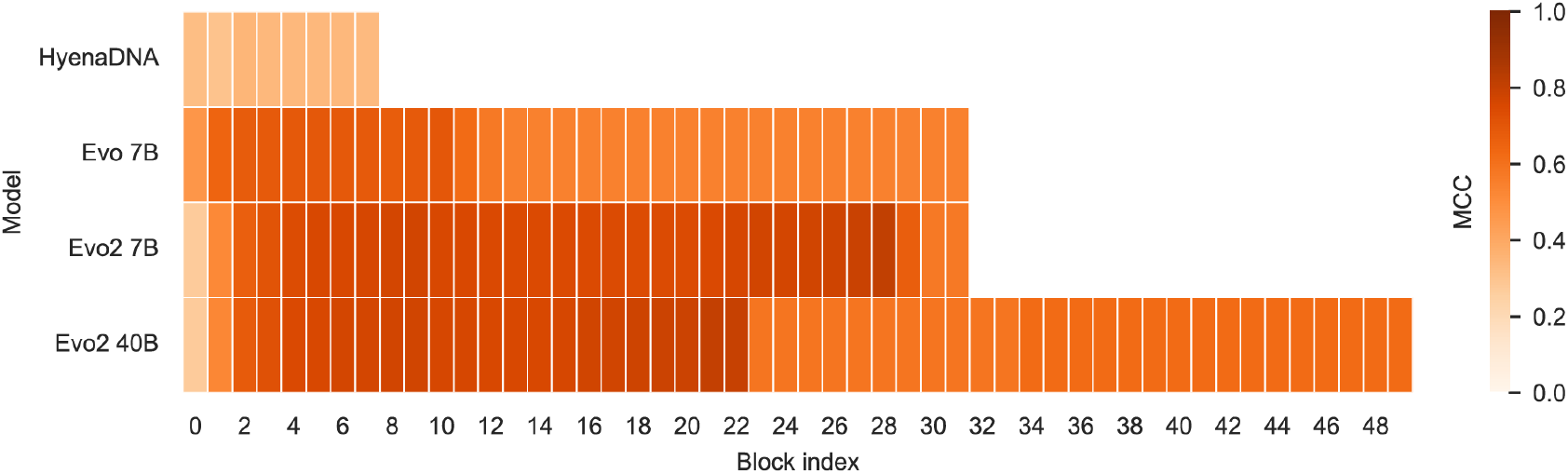
Results of the CDS annotation. These results show the predictive performance of embedding-based logistic regression classifiers on the annotation dataset.

### Logit lens analysis of autoregressive gLMs

To obtain a deeper understanding of autoregressive gLMs, we applied the logit lens to the long-context autoregressive gLMs: HyenaDNA, Evo and Evo 2 (7B and 40B). We identified saturation of the NLL before the final layer in both the Evo and Evo 2 models (Figure 5A and Supplementary Figure S8). The decrease in NLL at intermediate layers became particularly prominent in Evo and Evo 2 (40B), where more than half of the deeper layers presented NLL values comparable to the model outputs. Remarkably, the onset of NLL reduction was accompanied by a marked increase in the L2 norm of the embedding representations, suggesting a major change in the scale of latent representation during the later stages of processing. Moreover, for these gLMs, the decrease in NLL observed in deeper layers coincided with a decrease in downstream task performance (Figure 5B and Supplementary Figure S8).

**Figure 5.**
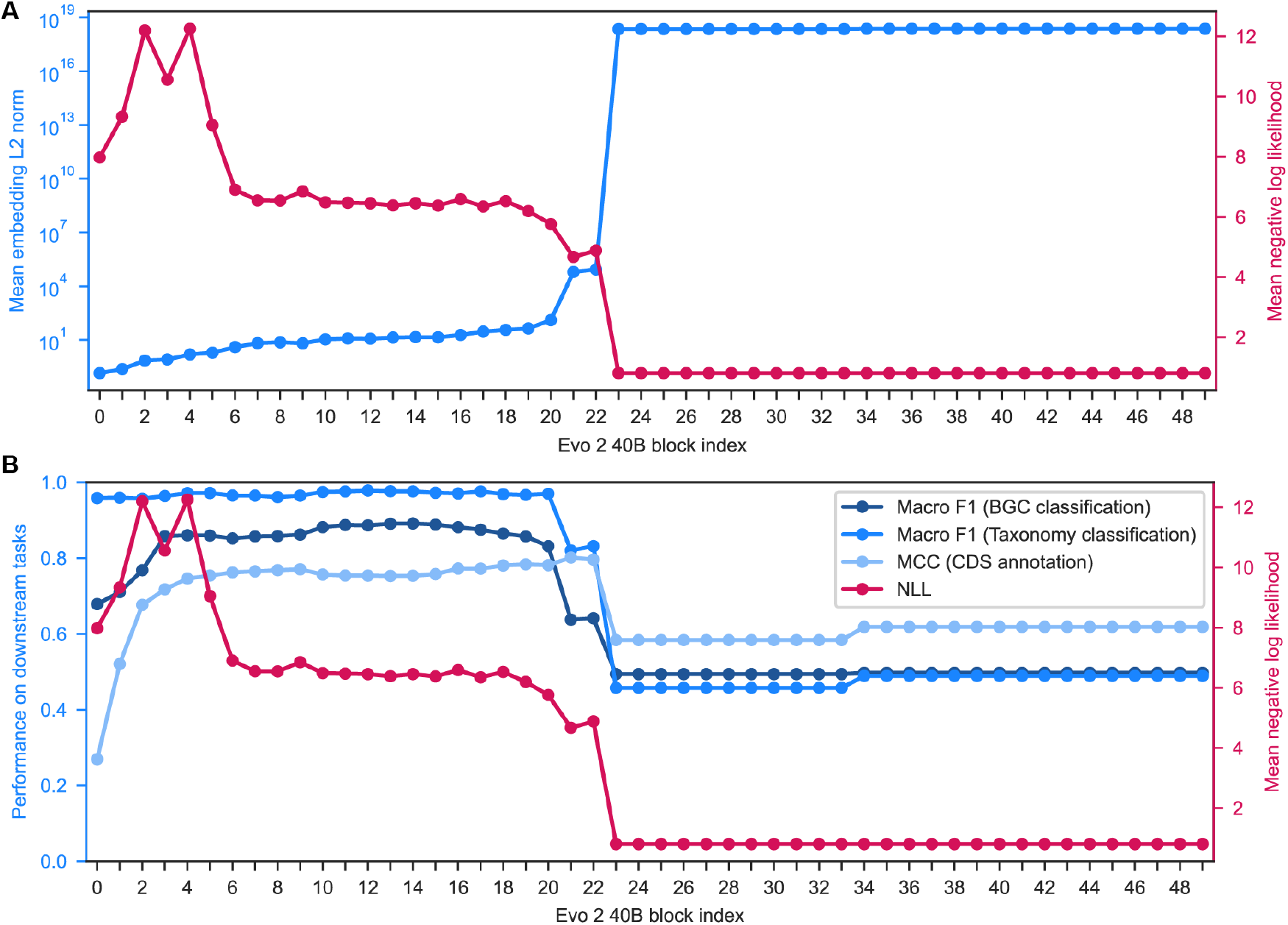
Logit lens to Evo 2 (40B). (A) Relationships between the NLL and the L2 norm of Evo 2 (40B) embeddings. (B) Relationships between the NLL and the downstream task performance for Evo 2 (40B).

## Discussion

Recent long-context gLMs represent breakthroughs in genomics, but the understanding of the properties of gLMs remains limited. To address this issue, we introduced BGCs-Bench, a unified benchmark focused on BGCs for assessing long-sequence modeling. We conducted a systematic and layer-wise evaluation of existing long-context gLMs using BGCs-Bench, which enabled characterization of the embeddings across models, depth and tasks.

The benchmarking results revealed that long-context gLMs except for HyenaDNA could yield embeddings that were more effective than k-mer frequency for downstream tasks when appropriate intermediate layers were chosen. These findings underscore the importance of layer selection in extracting high-quality representations from long-context gLMs. In contrast, the relatively weak performance of HyenaDNA may be attributable to a taxonomic mismatch between its pretraining data and BGCs-Bench. While HyenaDNA was pretrained using solely the human genome, our benchmarking datasets comprise mainly bacterial and fungal sequences.

Previous studies on NLP and protein language models have demonstrated that language models capture distinct information through different layers [22,23]. On the basis of these findings, we initially hypothesized that gLMs would likewise encode functional, phylogenetic and syntactic signals across model depth such that the best embedding representations would vary for downstream tasks. However, our evaluation of gLM embeddings on diverse downstream tasks suggested that signals relevant to different levels of biological meaning appeared to be captured collectively within some intermediate layers in the Evo and Evo 2 models. That is, distinct biological signals were not clearly partitioned across layers in the StripedHyena-based gLMs.

In addition, the logit lens to autoregressive gLMs revealed that NLL saturated early in Evo and Evo 2 (40B), raising the possibility that these models are parameter-inefficient. In StripedHyena-based gLMs, the NLL approached the model output before the last layer, and this reduction in NLL coincided with a decrease in downstream task performance. To our knowledge, this is the first study to characterize gLM embeddings by integrating downstream task evaluation with the logit lens technique. Together with the observation in the layer-wise assessment of embedding quality for downstream tasks, this pattern suggests that StripedHyena-based gLMs may contain separable functional regimes across depth: intermediate layers to capture biologically meaningful information from input DNA sequences, and deeper layers to optimize latent representations for sequence generation. Provided that the decrease in NLL was accompanied by an increase in the L2 norm of the embeddings, these gLMs improve the pretraining objective by inflating logits and thereby sharpening the probability distribution after a softmax function. One possible interpretation of these results is that next-token prediction over the four-letter nucleotides may be too simple a pretraining objective for an LLM architecture. Moreover, given that a similar decay in the contribution of deeper layers, termed the “Curse of Depth” [30], has been commonly observed in various LLMs, this phenomenon may reflect a general property of large-scale autoregressive models.

A limitation of this study is that the upper bound of the DNA sequence length in the datasets was restricted to 131 kbp or 66 kbp because of the maximum context lengths of existing gLMs and computational constraints. However, this operation removed at most 16% of the original data, and more than 60% of the excluded sequences were NRPS or PKS BGCs, which were abundant in the original source dataset. Taken together, the influence of this filtering is expected to be limited.

## Conclusion

In summary, we introduce BGCs-Bench, a unified benchmark for assessing long-context genomic modeling with a focus on BGCs. For existing long-context gLMs, we performed systematic and layer-wise evaluations of embedding representations, demonstrating that layer selection is a key factor for downstream tasks. Furthermore, the logit lens analysis suggests that state-of-the-art autoregressive gLMs are likely to overfit the pretraining task. These insights will offer communities the opportunity for more effective development and application of long-context gLMs. Overall, BGCs-Bench establishes a valuable resource for future efforts towards biologically meaningful genome modeling.

## Key points

▪ BGCs-Bench is a unified benchmark focused on biosynthetic gene clusters (BGCs) for assessing long-sequence modeling that includes three complementary downstream tasks: biosynthetic class prediction, taxonomic classification, and CDS annotation.
▪ Systematic layer-wise evaluation of long-context gLMs on BGCs-Bench reveals that foundational gLMs can capture biological signals at various levels, whereas layer selection is important for successes in downstream tasks.
▪ Together with the results of downstream tasks, logit lens suggests that StripedHyena-based gLMs consist of earlier layers to encode biologically meaningful information from inputs and deeper layers to optimize embeddings for sequence generation.

## Acknowledgement

Computations were partially performed on the NIG supercomputer at ROIS National Institute of Genetics.

## Author contributions

Keisuke Hirota (Conceptualization, Formal Analysis, Investigation Resources, Software, Writing—original draft), Koichi Higashi (Conceptualization, Supervision, Writing—review & editing), Ken Kurokawa (Conceptualization, Funding acquisition, Supervision, Writing—review &editing), and Takuji Yamada (Conceptualization, Funding acquisition, Supervision, Writing— review & editing).

## Data availability

Our benchmark datasets and codes are available at https://github.com/GenAIBio/BGCs-Bench.

