## supplementary_figures for "Benchmarking long-context genome language models on biosynthetic gene clusters"

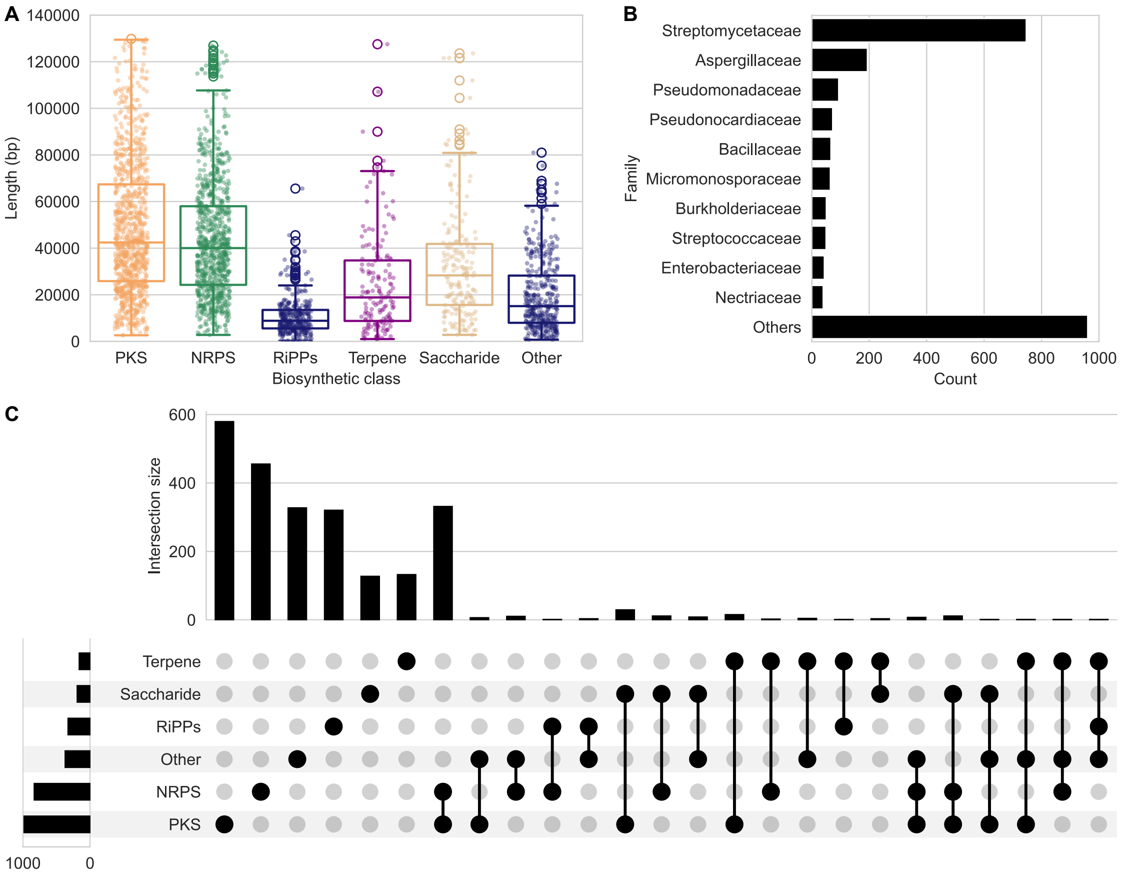


**Supplementary Figure S1. Details of the 131k dataset.** (A) Sequence length distribution in the 131k dataset. (B) Family distribution in the 131k dataset. (C) Biosynthetic class distribution in the 131k dataset.

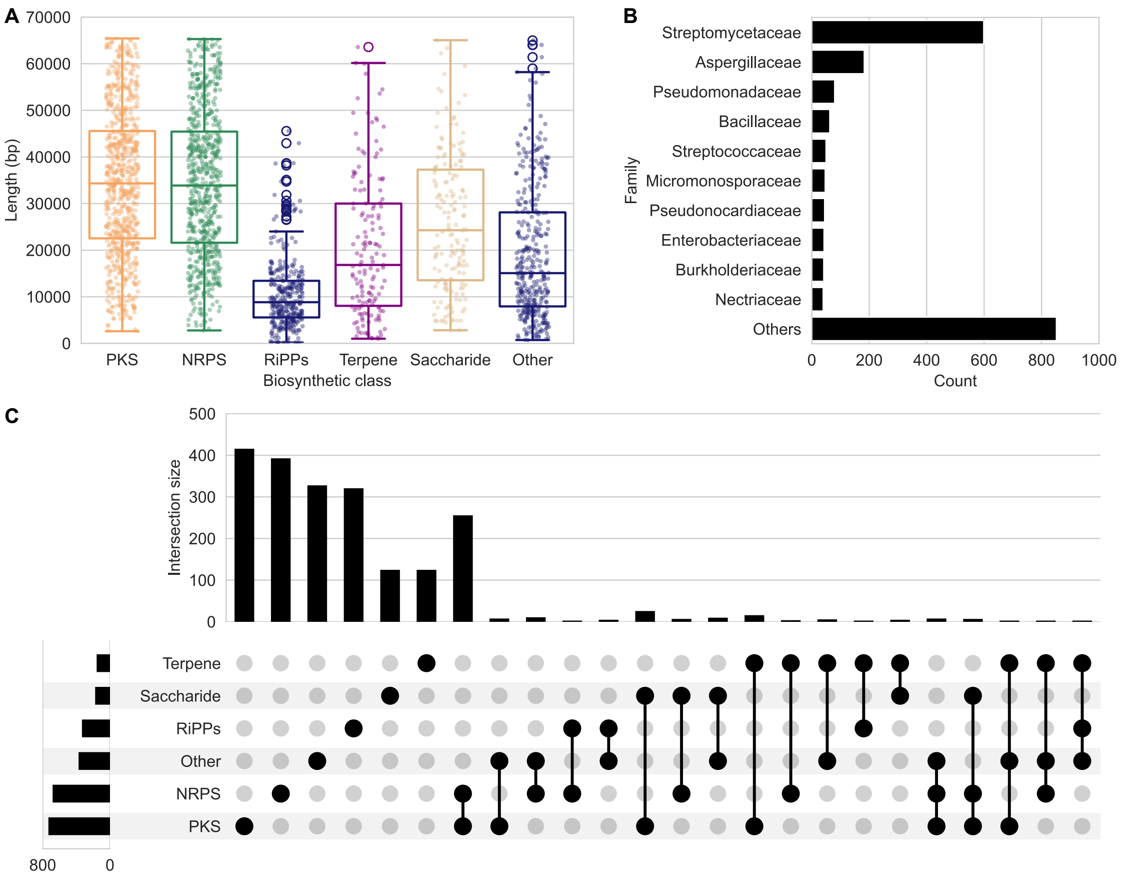


**Supplementary Figure S2. Details of the 66k dataset.** (A) Sequence length distribution in the 66k dataset. (B) Family distribution in the 66k dataset. (C) Biosynthetic class distribution in the 66k dataset.

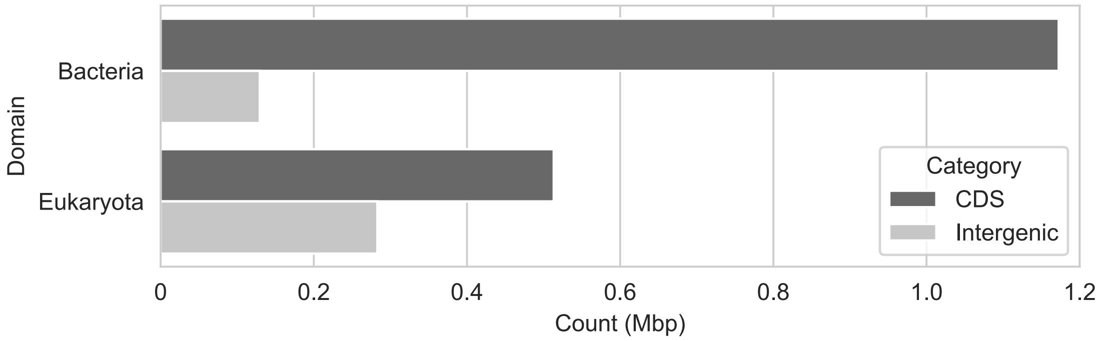


**Supplementary Figure S3. Counts of nucleotides in the annotation datasets.** Total nucleotides of CDSs and intergenic regions for bacteria and eukaryota in the annotation dataset.

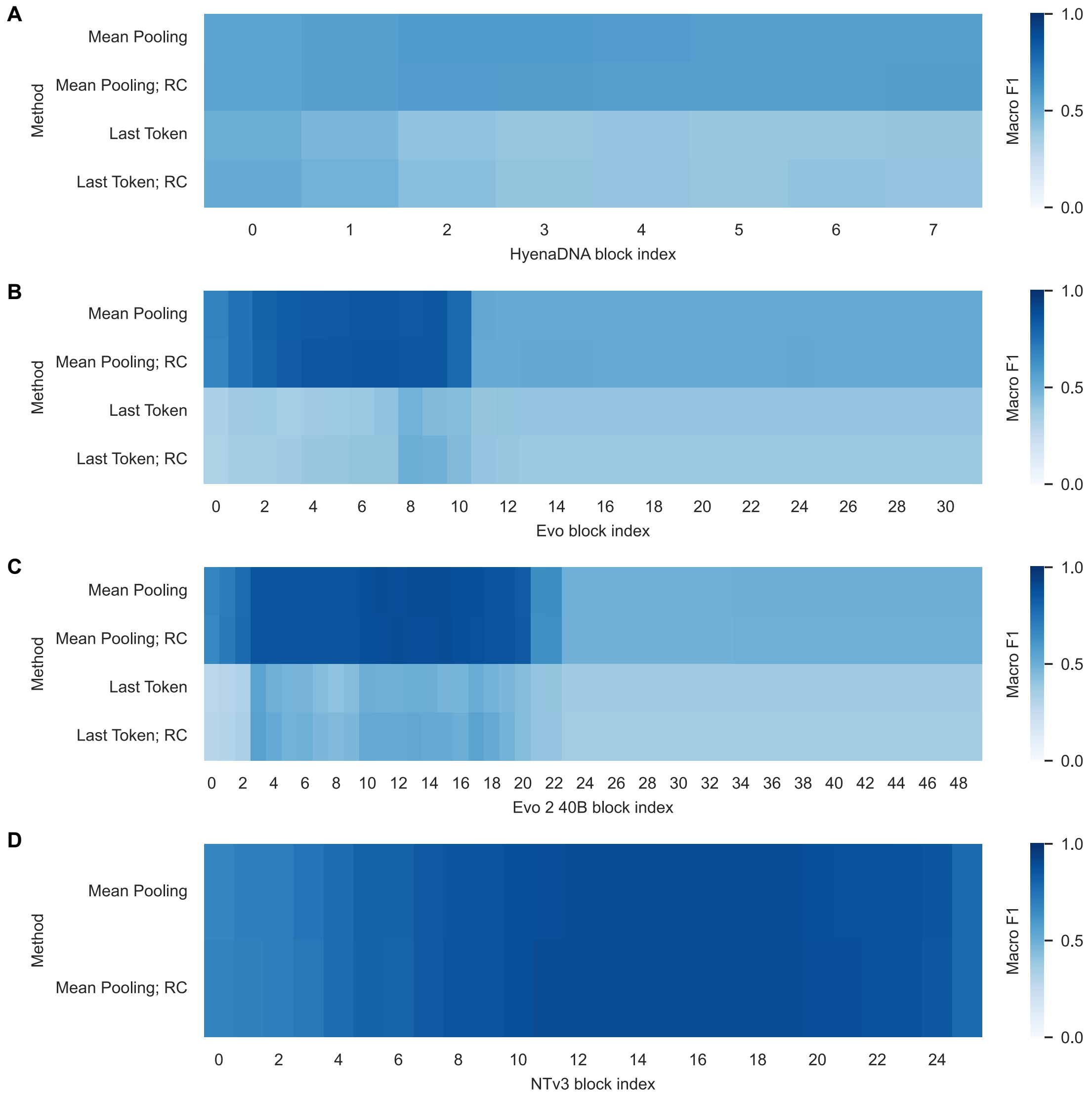


**Supplementary Figure S4. Results of the biosynthetic class prediction for each gLM.** These results show the predictive performance of embedding-based logistic regression classifiers on the 66k dataset excluding the "Other" class. (A) Performance of HyenaDNA-based classifiers. (B) Performance of Evo-based classifiers. (C) Performance of Evo 2 (40B)-based classifiers. (D) Performance of NTv3-based classifiers.

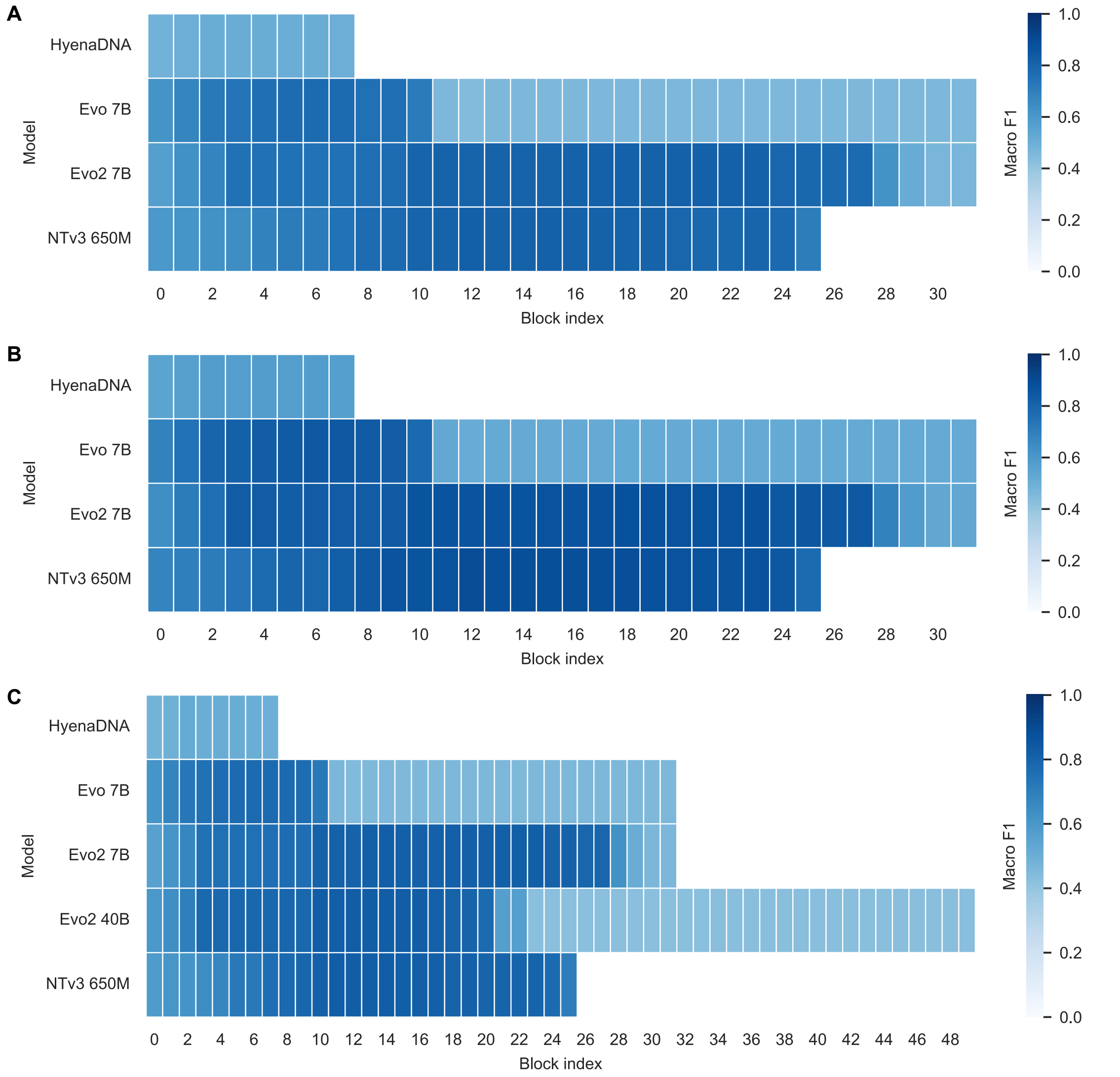


**Supplementary Figure S5. Results of the biosynthetic class prediction for each setting.** (A) Overview of the predictive performance per layer in the gLMs on the 131k dataset including the "Other" class. (B) Overview of the predictive performance per layer in the gLMs on the 131k dataset excluding the "Other" class. (C) Overview of the predictive performance per layer in the gLMs on the 66k dataset including the "Other" class.

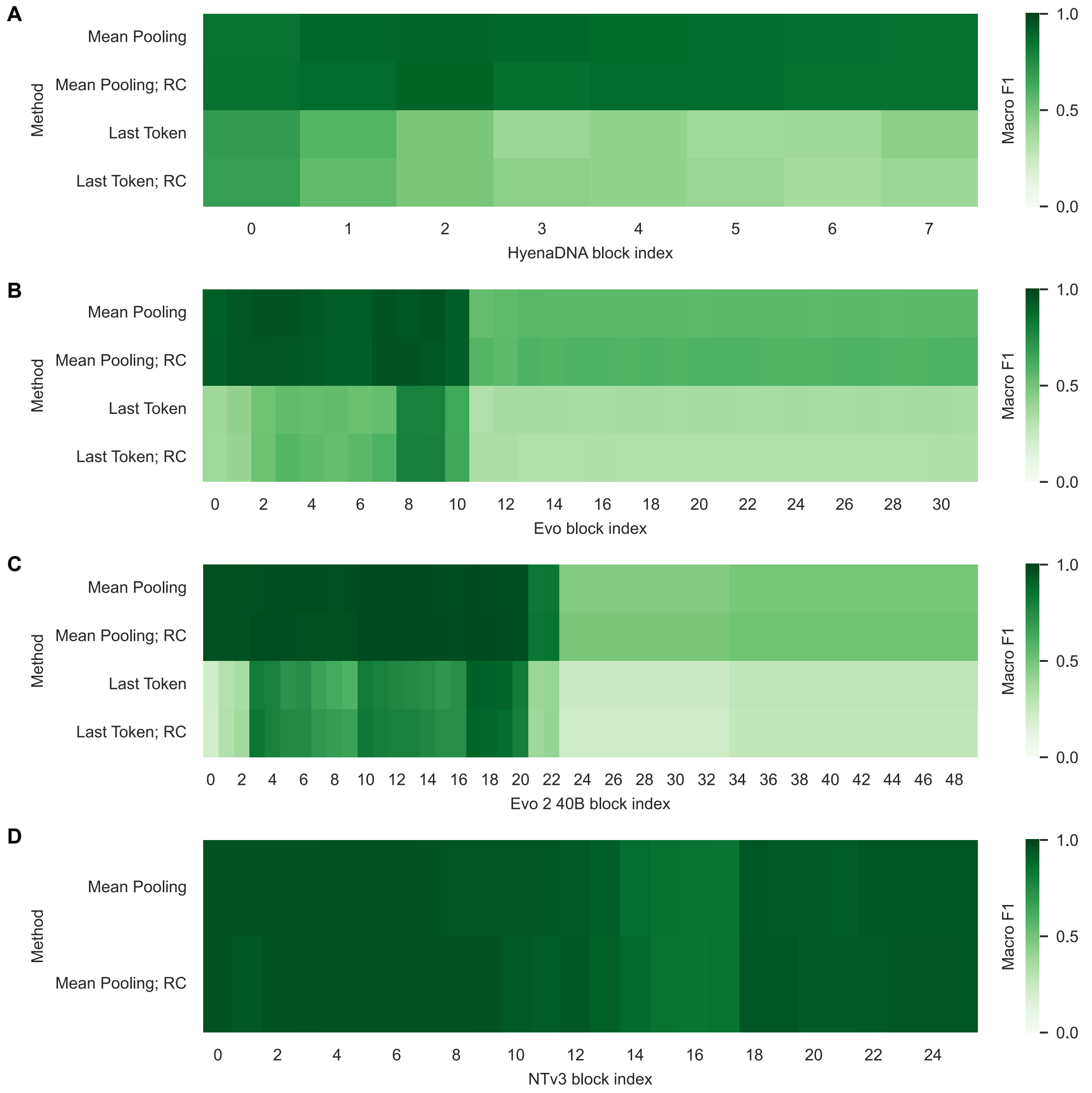


**Supplementary Figure S6. Results of the taxonomic classification for each gLM.** These results show the predictive performance of embedding-based logistic regression classifiers on the 66k dataset excluding "Other" families. (A) Performance of HyenaDNA-based classifiers. (B) Performance of Evo-based classifiers. (C) Performance of Evo 2 (40B)-based classifiers. (D) Performance of NTv3-based classifiers.

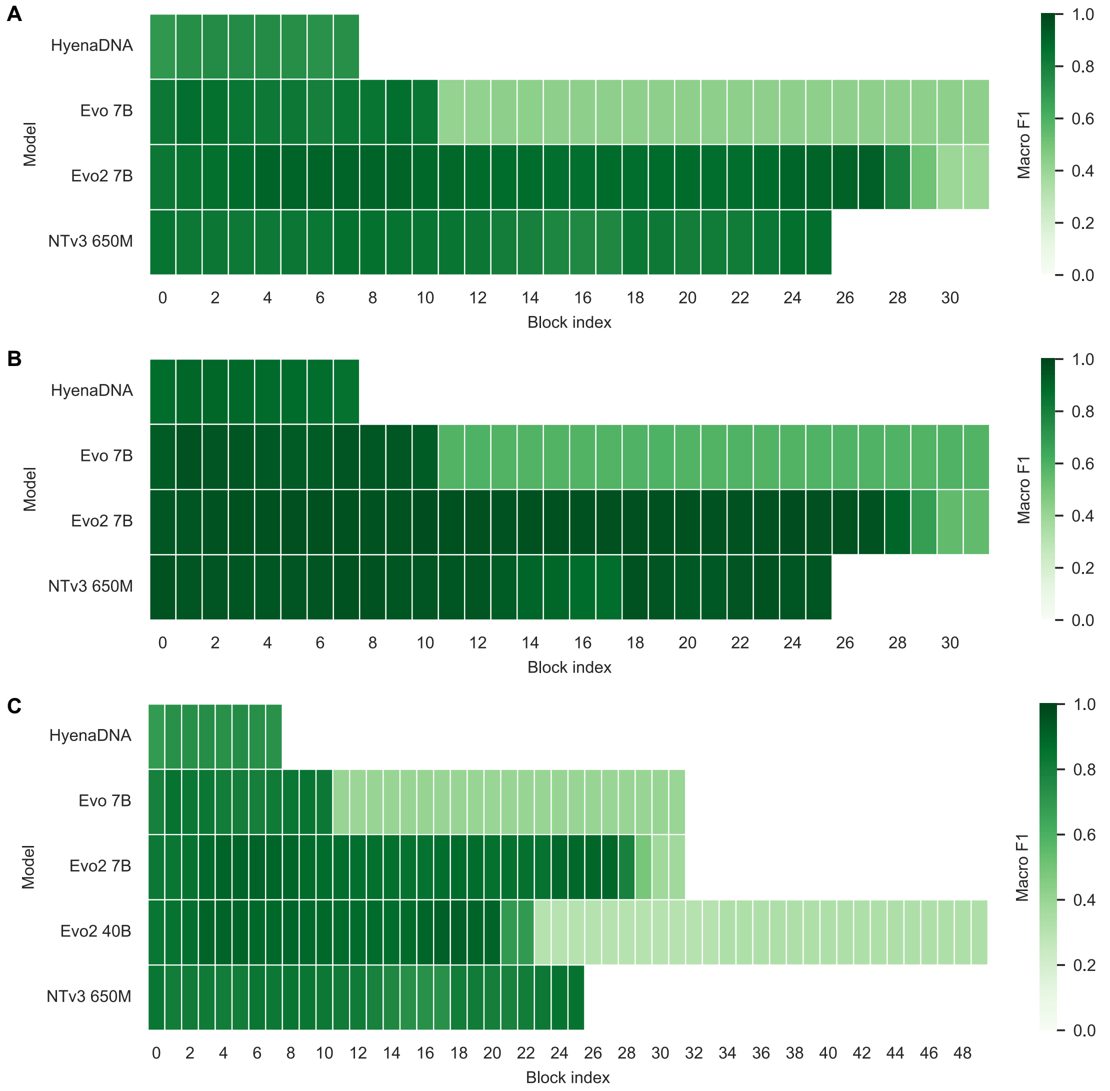


**Supplementary Figure S7. Results of the taxonomic classification for each setting.** (A) Overview of the predictive performance per layer in the gLMs on the 131k dataset including "Other" families. (B) Overview of the predictive performance per layer in the gLMs on the 131k dataset excluding "Other" families. (C) Overview of the predictive performance per layer in the gLMs on the 66k dataset including "Other" families.

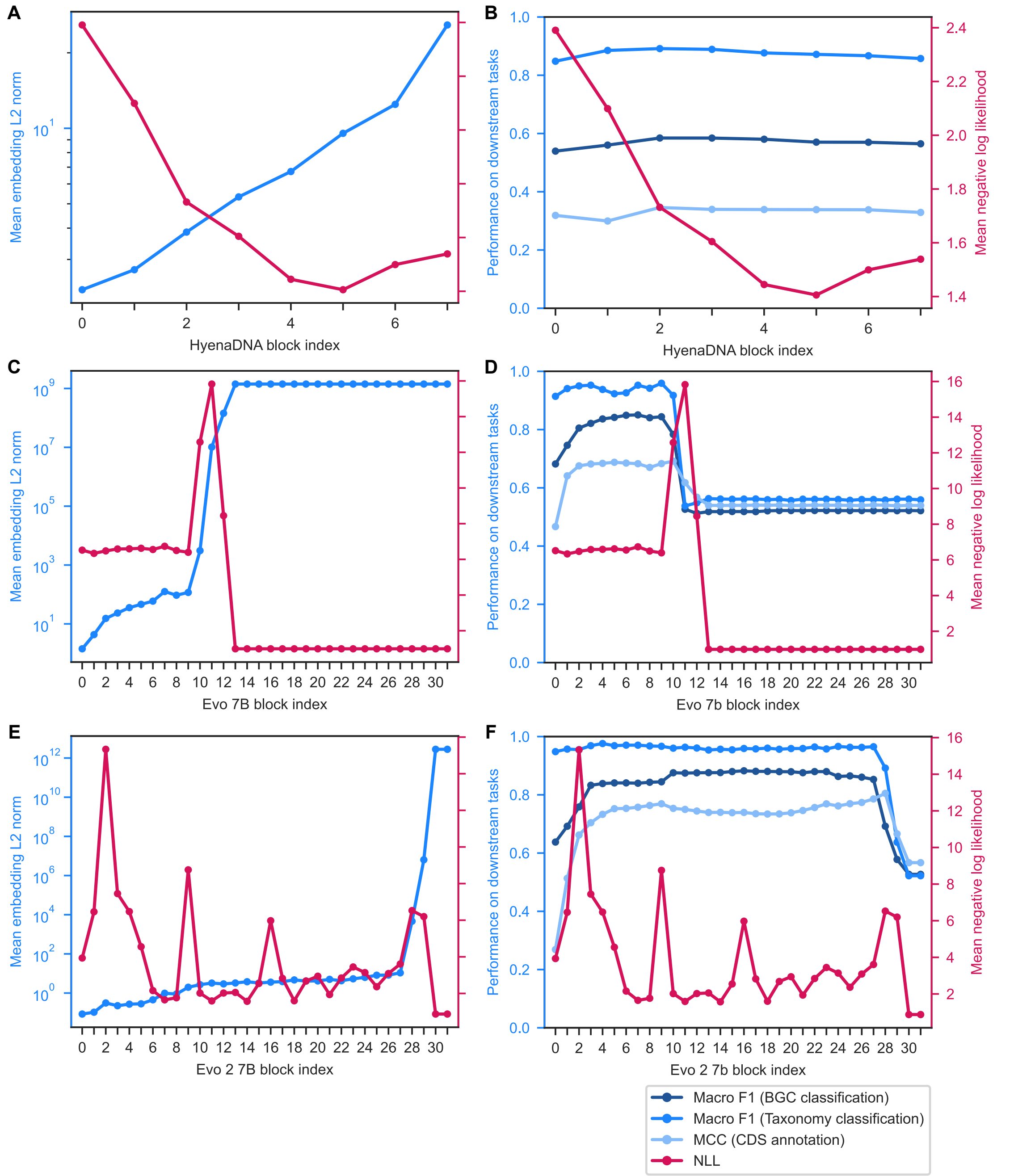


**Supplementary Figure S8. Logit lens to autoregressive gLMs.** (A) Relationships of the NLL and L2 norm of HyenaDNA embeddings. (B) Relationships of the NLL and the downstream task performance for HyenaDNA. (C) Relationships of the NLL and L2 norm of Evo embeddings. (D) Relationships of NLL and the downstream task performance for Evo. (E) Relationships of the NLL and L2 norm of Evo 2 (7B) embeddings. (F) Relationships of the NLL and the downstream task performance for Evo 2 (7B).
